# Emerging SARS-CoV-2 mutation hotspots associated with clinical outcomes

**DOI:** 10.1101/2021.03.31.437666

**Authors:** Xianwu Pang, Pu Li, Lifeng Zhang, Lusheng Que, Min Dong, Qihui Wang, Yinfeng Wei, Bo Xie, Xing Xie, Lanxiang Li, Chunyue Yin, Liuchun Wei, Qingniao Zhou, Yingfang Li, Lei Yu, Weidong Li, Zengnan Mo, Jing Leng, Yanling Hu

**Author notes:** Corresponding author: Yanling Hu. These authors contributed equally (Xianwu Pang, Pu Li, Lifeng Zhang, Lusheng Que, Min Dong).

## Abstract

Severe acute respiratory syndrome coronavirus 2 (SARS-CoV-2) is the cause of the ongoing coronavirus disease 2019 (COVID-19) pandemic. Understanding the influence of mutations in the SARS-CoV-2 gene on clinical outcomes and related factors is critical for treatment and prevention. Here, we analyzed 209,551 high-coverage complete virus sequences and 321 RNA-seq samples to mine the mutations associated with clinical outcome in the SARS-CoV-2 genome. Several important hotspot variants were found to be associated with severe clinical outcomes. Q57H variant in ORF3a protein were found to be associated with higher mortality rate, and was high proportion in severe cases (39.36%) and 501Y.V2 strains (100%) but poorly proportional to asymptomatic cases (10.04%). T265I could change nsp2 structure and mitochondrial permeability, and evidently higher in severe cases (20.12%) and 501Y.V2 strains (100%) but lower in asymptomatic cases (1.43%). Additionally, R203K and G204R could decrease the flexibility and immunogenic property of N protein with high frequency among severe cases, VUI 202012/01 and 484K.V2 strains. Interestingly, the SARS-CoV-2 genome was more susceptible to mutation because of the high frequency of nt14408 mutation (which located in RNA polymerase) and the high expression levels of ADAR and APOBEC in severe clinical outcomes. In conclusion, several important mutation hotspots in the SARS-CoV-2 genome associated with clinical outcomes was found in our study, and that might correlate with different SARS-CoV-2 mortality rates.

## Introduction

Coronavirus infectious disease 2019 (COVID-19) is caused by the severe acute respiratory syndrome coronavirus 2 (SARS-CoV-2). COVID-19 has rapidly spread worldwide and became a global health emergency. The clinical spectrum of SARS-CoV-2 infection appears to be wide, encompassing asymptomatic infection, mild upper respiratory tract illness, and severe viral pneumonia that may result in respiratory failure and even death, with many patients being hospitalized with pneumonia[1-3]. According to the World Health Organization, the mortality rate as of October 2020 is about 2.7%. However, the mortality rate is up to 26% in severe cases[4]. The ongoing rapid spread of the virus worldwide, coupled with asymptomatic cases and patients with severe symptoms, raises an important concern that it may further mutate into more highly transmissible or virulent forms.

Mortality rates can widely vary according to geography, demographics and healthcare infrastructure[5], as well as according to various host factors, including advanced age, being male, and comorbidities[6, 7]. However, the viral factors underlying the severity of COVID-19 disease and the corresponding mortality rate remain unclear. Some mutations in the SARS-CoV-2 genome can reportedly remarkably change the virus’ properties, such as transmission modes and rates, as well as the ability to cause diseases[8-10]. The G614D variant in the S glycoprotein is associated with increased transmissibility, infectivity, and viral loads but not with disease severity[9, 11]. Parinita et al[12] suggested that the presence of P25L in ORF3a is a probable mechanism of immune evasion and likely contributes to enhanced virulence, which are associated with higher mortality rates in SARS-CoV-2 infection. Similarly, some mutations, such as S197L, S194L, and a 382-nucleotide deletion, are associated with clinical outcomes[13, 14]. Additionally, mutation types, such as the G/T variant in the open reading frame 1ab (ORF1ab) gene, is associated with clinical infections[15].

Therefore, RNA virus mutations not only contributes to viral adaptation that creates a balance between the integrity of genetic information and genome variability[16-18] but also lead to different clinical outcomes. Understanding the characteristics of viral mutations is essential for the pathogenesis, immune escape, drug resistance, vaccine design, antiviral therapy, and diagnosis of COVID-19. In this context, we focused on SARS-CoV-2 mutations associated with clinical outcomes. First, 209,551 high-coverage complete virus sequences from the GISAID database were compared with the SARS-CoV-2 reference genome (NC_045512.2). This study aimed to gain important insights into virus mutations and their occurrence over time, especially hotspot mutations associated with different clinical outcomes. Second, RNA-dependent RNA polymerase (RdRp) and RNA-editing enzyme analyzed from 321 RNA-seq samples were explored to explore the SARS-CoV-2 mutations that are associated with different clinical outcomes.

## MATERIALS AND METHODS

### Data source

Publicly available SARS-CoV-2 complete genomes of different patients were collected from the GISAID virus repository (https://www.gisaid.org/) from January 1, 2020 to January 1, 2021. Viral sequences with a complete genome (28000–30,000 bps) were included in this study. Moreover, the viral genomic sequences of MERS and SARS were downloaded from National Center for Biotechnology Information (NCBI; https://www.ncbi.nlm.nih). RNA-seq datasets available from the PRJNA601736 (2 sample, Raw sequence reads), PRJNA603194 (1 sample, Raw sequence reads), PRJNA605907 (8 sample, Raw sequence reads), PRJNA631753 (62 sample, Raw sequence reads), PRJNA639791 (6 sample, Raw sequence reads), PRJNA656568 (93 SARS-CoV-2 positive samples, 99 samples without any virus infection, Transcriptome or Gene expression) and PRJNA683226 (37 mild samples, 10 moderate samples, 3 severe samples, Transcriptome or Gene expression) projects were also downloaded from the NCBI (https://www.ncbi.nlm.nih.gov/sra/).

### Single nucleotide variation (SNV) calling in genomic sequence

SNVs were called with a custom R script, by comparing the viral genome sequences to the reference sequence. The reference of SARS-CoV-2 is NC_045512.2 sequence, that of SARS is NC_004718.3 sequence, and that of MERS is NC_019843.3 sequence. SNVs occurring on coding sequences were annotated with custom R scripts to determine the outcome of nucleotide changes, including nonsense, nonsynonymous, and synonymous mutations.

### SNV calling in the transcriptome of SARS-CoV-2

REDItools 2[19, 20] and JACUSA[21] were used together to call the SNVs in the transcriptome of SARS-CoV-2. With REDItools 2, all SNVs within 15 nucleotides from the beginning or the end of the reads were removed to avoid artifacts due to misalignments. Furthermore, the AS_StrandOddsRatio parameter was utilized to avoid potential artifacts due to strand bias, and any mutation with an AS_StrandOddsRatio of > 4 was removed from the dataset. The overlap of mutations generated by REDItools 2 and JACUSA were considered. The threshold to filter the SNVs was based on minimum coverage (20 reads), number of supporting reads (at least four mutated reads), allelic fraction (0.5%), quality of the mapped reads (>25), and base quality (>35).

### Gene expression in transcriptome

Gene expression in transcriptome was analyzed by introducing transcript per million (TPM) to estimate gene expression levels. Given that TPM considers the effects of sequencing depth, gene length, and sample on reads count, it is often used to estimate gene expression levels. In this study, all samples were processed through a SARS-CoV-2 reference-based assembly pipeline that involved removing non-SARS-CoV-2 reads with Kraken2[22] and aligning to the SARS-CoV-2 reference genome NC_045512.2 by using Samtools[23]. SARS-CoV-2 TPM was calculated using the R package tximport[24] via the length Scaled TPM method.

### Statistical analysis

The normality of data distribution was checked via Shapir–Wilk test. Categorical variables were expressed as absolute frequency and percentages. The different of SNV counts per genome between different clinic outcome was analyzed using one-way ANOVA followed by Student Newman–Keuls test, and Chi-square was utilized to compare the frequency of mutation sites between different clinical outcomes. Wald test and log2 FDR were used to compare the number of gene expression between different groups. Pearson correlation coefficient was calculated to estimate the correlation between sample gene expression and viral load. A higher Pearson correlation coefficient indicated a stronger correlation. All p-values were calculated from two-sided tests using 0.05 as the significance level.

## Results

### Characterization of mutations in the SARS-CoV-2 genome

A total of 209,551 complete SARS-CoV-2 genome sequences isolated from patients were download from the GISAID Database. All sequences were aligned and compared with the SARS-CoV-2 reference genome (NC_045512.2). Results showed that C>T, G>T, A>G and G>A mutations were the main SNV types (Figure 1A), and the frequency of these SNV types increased, especially the C>T mutation from January 2020 to January 2021 (Figure 1B). The effects of clade, region, sex, and age on base mutations were also analyzed, indicating that clade and region were also the main factors of mutation. For example, GV was more prone to C>T mutations, while GR was more prone to A>G, G>A and G>T mutations. Furthermore, Europe and Africa were more prone to all base mutations than the other regions (Figure S1). The sequence context of mutation showed special preference, the C>T mutation preferentially targets a 5□-TTCTA-3 □ motif, the A>G mutation preferentially targets a 5□-GGATG-3□ motif, and the G>A mutation preferentially targets a 5 □-AAGGG-3 □ motif (Figure S2).

**Figure 1.**
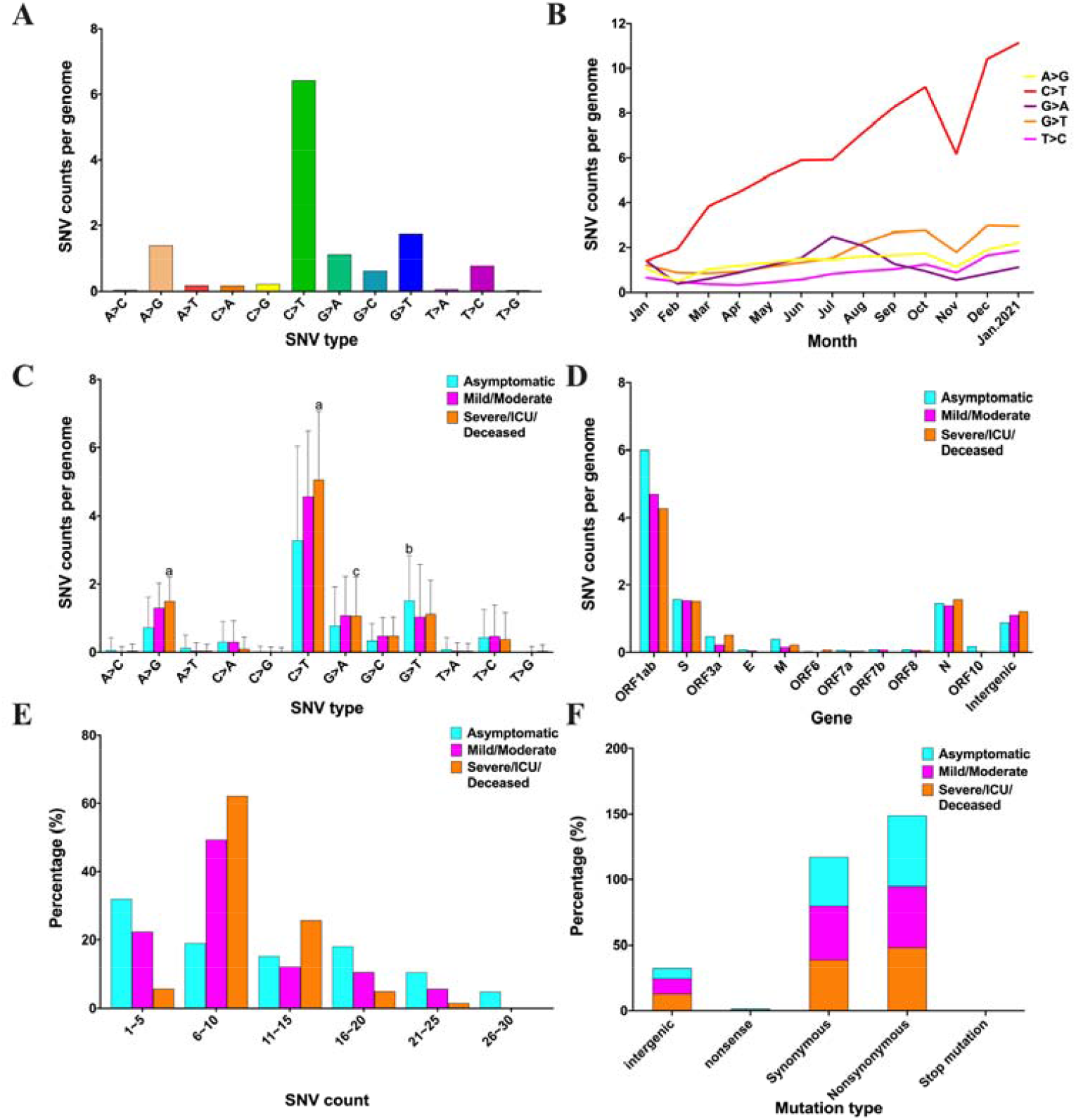
Characterization of mutations in the SARS-CoV-2 genome. A: SNV type in the SARS-CoV-2 genome. B: Dynamics of SNV counts over time. C: Number of different SNV types distributed in the three clinical outcomes. D: SNVs of three clinical outcomes distributed in the gene of SARS-CoV-2. E: Ranges of SNV counts in the three clinical outcomes. F: Mutation results of three clinical outcomes. ^a^ *p* < 0.001, ^b^ *p* < 0.01, ^c^ *p* < 0.05 represents there is significant difference among the three clinical outcomes. All results were analyzed using one-way ANOVA followed by Student Newman-Keuls test.

A total of 1,329 high-quality sequences had special clinical information. According to clinical description, these sequences were divided into three groups: asymptomatic, mild/moderate, and severe/ICU/deceased. The mutation frequency of C>T, A>G, and G>A was substantially higher in severe cases than in asymptomatic cases, whereas that of G>T was higher in asymptomatic case than that in other groups (Figure 1C). These mutations were concentrated in the ORF1ab, S, N genes (Figure 1D). The count of SNV per genome was divided into six groups: asymptomatic cases were mainly distributed in 1–5 (32%) and 6–10 (19.1%), mild/moderate cases were largely distributed in 1–5 (22.3%) and 6–10 (49.4%), and severe cases were chiefly distributed in 6–10 (62.2%) and 11–15 (25.8%). The difference in the three clinical outcomes was statistically significant (*p*<0.000) (Figure 1E). The mutation results of asymptomatic, mild/moderate, and severe cases were primarily nonsynonymous (54%, 46.5%, and 48.3%, respectively), followed by synonymous (37%, 41.1%, and 38.7%, respectively) (Figure 1F). The sequence context of mutation showed that the motif preference for the three clinical outcomes was similar (Figure S3).

### Identification of mutation hotspots in different clinic outcomes

A total of 2,093 mutation sites were found in the three clinical outcomes (Figure S4). According to the results of the Chi-square test, 130 missense mutation sites were notably different in the three clinical outcomes (Table S1). These missense mutations were primarily distributed in ORF1ab (56.98%), S (16.28%), ORF3a (11.63%), and N (10.47%) (Figure 2A). With regard to the mutations that presented in over 5% of the samples, six missense mutations were present in asymptomatic cases only, including K4576N and N5542D in ORF1ab, A222V and S477N in S, A376T in N, and V30L in ORF10. Additionally, L3930F in ORF1ab and V1176F in S were found in severe symptom cases only (Figure 2B). Six major missense mutations were obviously higher in symptomatic cases than in asymptomatic cases: T265I in ORF1ab (asymptomatic:1%, symptomatic:15%), D614G in S (asymptomatic:59%, symptomatic:91%), Q57H in ORF3a (asymptomatic:10%, symptomatic:29%), R203K in N (asymptomatic:27%, symptomatic:40.4%), and G204R in N (asymptomatic:25%, symptomatic:40.4%) (Figure 2C).

**Figure 2.**
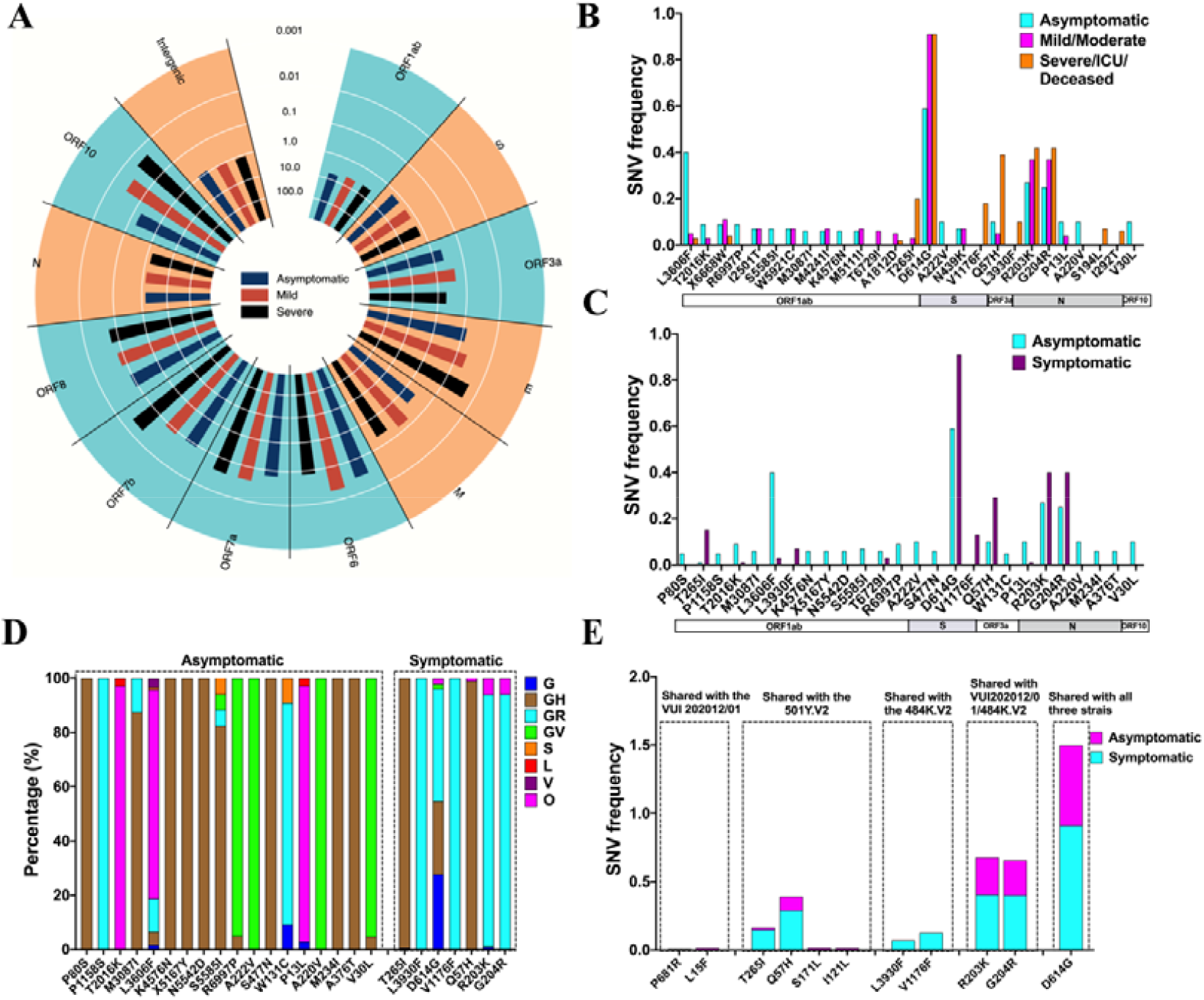
Identification of mutation hotspots in different clinical outcomes. A: Missense mutations of the three clinical outcomes distributed in different genes; the bar presents the incidence rate of missense mutation. B: Major SNV frequency in asymptomatic, mild/moderate, and severe outcomes. C: Major SNV frequency in asymptomatic and symptomatic outcomes. D: Mutation hotspots of asymptomatic and symptomatic outcome in clades. E: Major missense mutations occurring in asymptomatic and symptomatic cases shared with the SARS-CoV-2 VUI 202012/01, 501Y.V2, and 484K.V2 strains.

The mutations were found to be related to clades. Thus, the distribution of mutation hotspots among the clades were analyzed. In this study, 25 major missense mutation hotspots (frequency >5% in at least one of clade) were included in the analysis of distribution in different clades. For these mutations, 18 mutation hotspots were higher in the asymptomatic cases than in symptomatic cases: P80S, M3087I, K4576N, X5167Y, N5542D, S5585I, S477N, M220V, A376T, M234I, and A376T were predominantly distributed in GH clade; R6997P, A222V, A220V, and V30L were mainly distributed in GV clade; T2016K, L3606F, and P13L were mostly distributed in O clade. However, seven mutation hotspots were higher in the symptomatic cases than in asymptomatic cases; L3930F V1176F, R203K, and G204R were chiefly distributed in GR clade; T265I and Q57H were mostly distributed in GH clade (Figure 2E). The issue of whether these mutation hotspots were also distributed in the three major pandemic strains, namely, SARS-CoV-2 VUI 202012/01 strains circulating in the United Kingdom, 501Y.V2 strains spreading in South Africa, and 484K.V2 strains dispersing in Brazil, was also analyzed: P681R and L15F were found to be distributed in the SARS-CoV-2 VUI 202012/01 strains. S171L and I121L were observed to be distributed in the 501Y.V2 strains. T265I and Q57H were determined to be distributed in 501Y.V2 strains. L3930F and V1176F were uncovered to be distributed in 484K.V2 strains. R203K and G204R were discovered to be distributed in the SARS-CoV-2 VUI202012/01strains and 484K.V2 strains. D614G was established to be distributed in the three strains (Figure 2F).

### Dynamics distribution of clinical outcome-associated mutations over time

The dynamics distribution of the 25 major mutation hotspots over time was further analyzed. A222V, A220V, R6997P, and V30L co-occurred and rapidly increased starting from July 2020. M3087I, X5167Y, K4576N, N5542D, A376T, and S5585I co-occurred and gradually increased. S477N started to rise in June but decreased again in July (Figures 3A and 3B). D614G increased from 74.6% to 99.9%. T265I, Q57H, R203K, and G204R started to increase in March and gradually decreased in July. However, all the mutations associated with symptomatic cases increased starting from November 2020 (Figures 3C and 3D).

**Figure 3.**
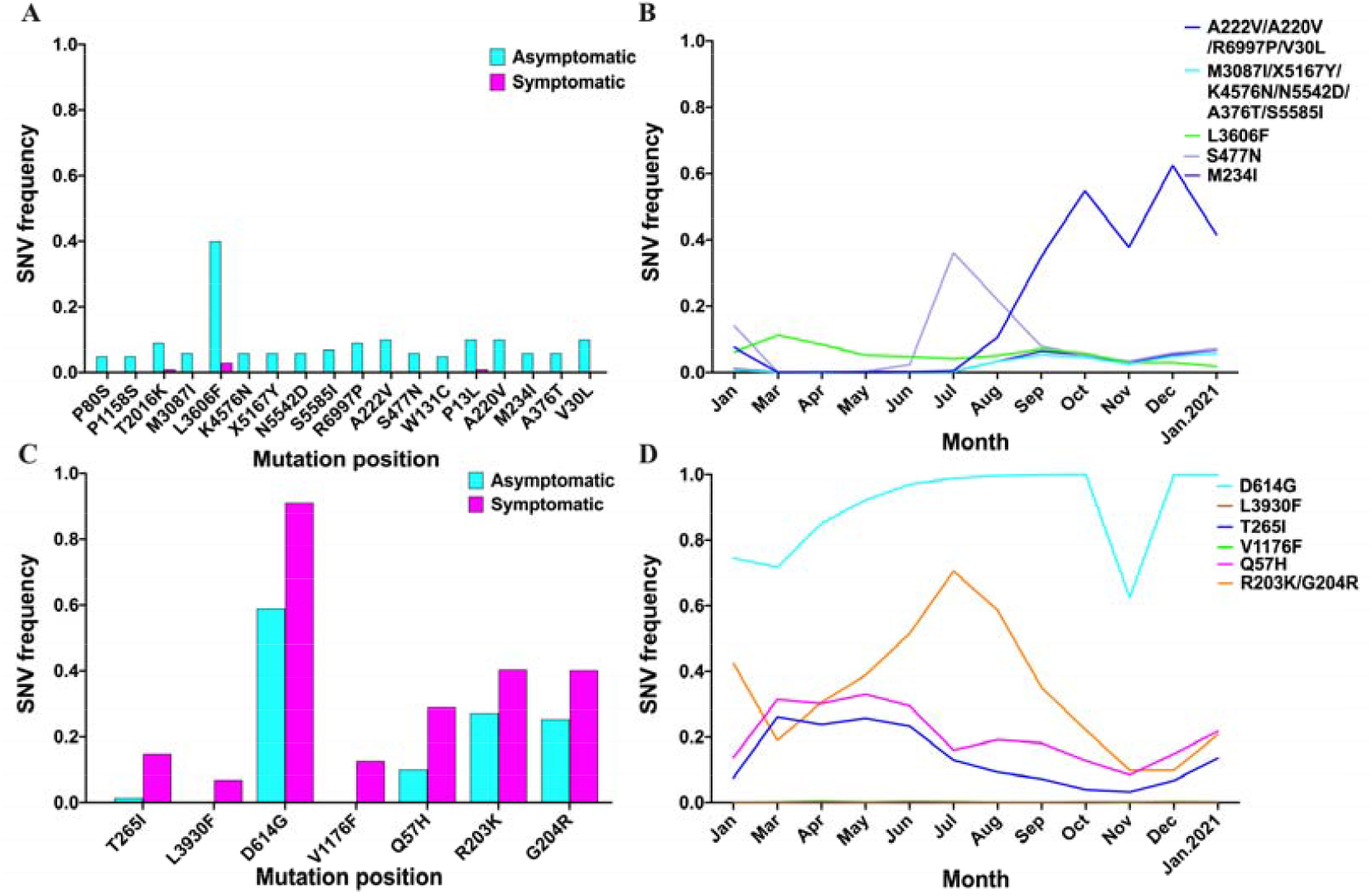
Dynamics distribution of major symptom-associated mutations. A: Major mutation sites in asymptomatic cases were higher than those in symptomatic cases, *p*< 0.05. B: Major SNV frequency of Figure 3A over time. C: Major mutation sites in symptomatic cases were higher than those in asymptomatic cases, *p*< 0.05. D: Major SNV frequency of Figure 3C over time.

### SNVs associated with RdRp mutations in different clinic outcomes

The issue of whether the mutation of RNA-dependent-RNA polymerase (RdRp) was associated with SARS-CoV-2 mutations was analyzed. Results showed that 142 mutations occurred in the RdRp. Moreover, the incidence rate of nt14408 mutation, which was distributed in asymptomatic cases (56.5%), mild/moderate cases (91.2%), and severe/ICU/deceased cases (95.9%), was the highest (Figure 4A). The 209,551 sequences were divided into “with nt 14408 mutation” and “without nt14408 mutation” groups, and the SNV count was calculated and divided into six ranges. According to the results of Chi-square test, 1–5 (*p*<0.000), 6–10 (*p*<0.000), 11–15 (*p*<0.000), 16–20 (*p*<0.000), and 21–25 (*p*=0.03) were significantly difference between “with nt 14408 mutation” and “without nt14408 mutation” groups, the “without nt14408 mutation” group was primarily distributed in 1–5, whereas the “with nt 14408 mutation” group was mostly distributed 6–10 and 11–15 (Figure 4B). The SNV type affected by nt14408 mutation was also analyzed. The counts of A>G, C>T, G>A, G>C and G>T in the “with nt14408 mutation” group were higher than those in the “without nt14408 mutation” group (Figure 4C).

**Figure 4.**
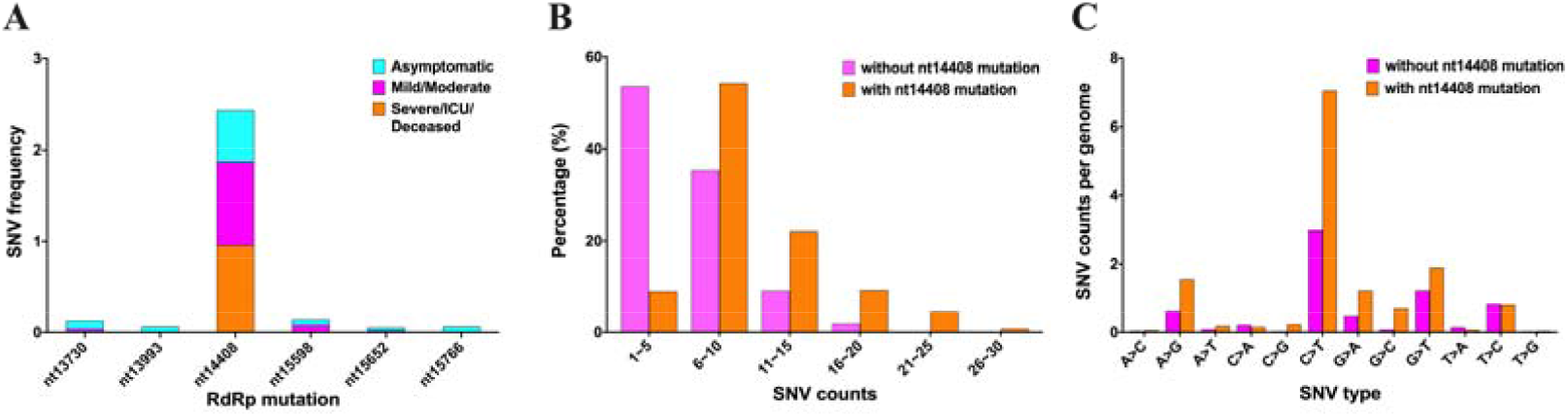
SNVs associated with RdRp mutation in different clinical outcomes. A: Major mutation sites of RdRp in the three clinical outcomes. B: Percentage of genome of different SNV count ranges in the “without nt14408 mutation” and “with nt14408 mutation” groups. C: SNV counts per genome of different SNV types in “without nt14408 mutation” and “with nt14408 mutation” groups.

### RNA editing in the transcriptome of SARS-CoV-2

The SNV type in the transcriptome of SARS-CoV-2 was analyzed to validate further the RNA editing in the body. These samples included lung tissues (Figure 5A), nasopharyngeal swabs (Figure 5C), and human alveolar type II cell organoids (Figure 5E). Finally, 19 lung tissue samples, 7 swab samples and 3 human alveolar type II cell organoids samples were qualified for further analysis. In each group, the distribution of all SNV types was different. C>T and G>T mutations were the major SNV types in lung tissues (Figure 5B), while A>G and T>C SNVs were the main SNV types in nasopharyngeal swabs (Figure 5 D), and A>G and C>T SNVs were the primary SNV types in human alveolar type II cell organoids (Figure 5F). The SNV types in every individual were also analyzed. The SNV types were different (Figure S5), which were pooled together (Figures 5, A, 5C and 5E).

**Figure 5.**
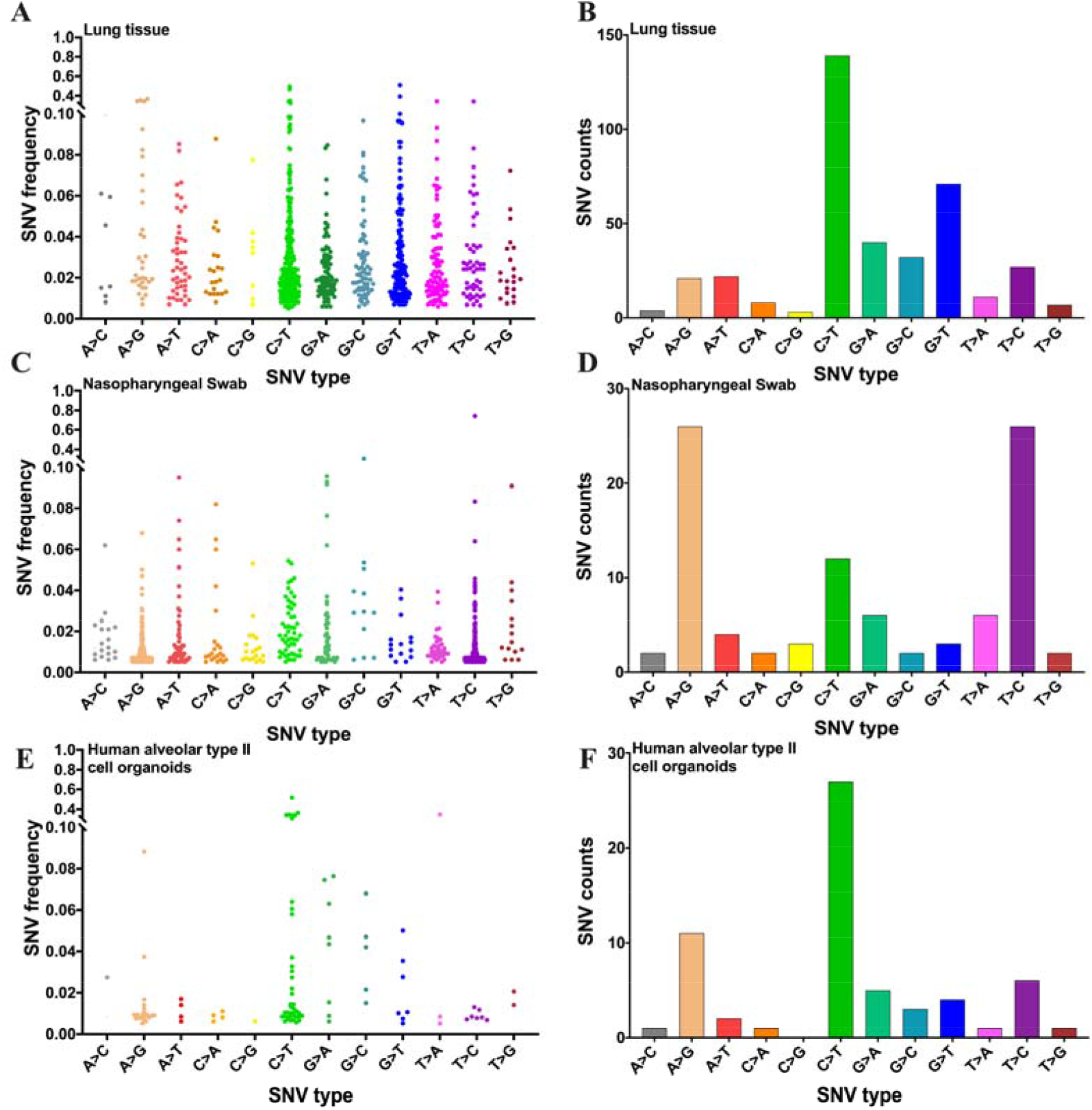
RNA mutation in the transcriptome of SARS-CoV-2. A: Distribution of all SNV types from lung tissues. B: Different SNV counts per sample from lung tissues. C: Distribution of all SNV types from nasopharyngeal swabs. D: Different SNV counts per sample from nasopharyngeal swabs. E: Distribution of all SNV types from human alveolar type II cell organoids. F: Different SNV counts per sample from human alveolar type II cell organoids.

### Expression of the RNA-editing enzyme in the transcriptome

In this study, PRJNA656568, PRJNA631753 and PRJNA683226 were used to analyze the expression of adenosine deaminase acting on RNA proteins (including ADAR1, ADARB1, and ADARB2), apolipoprotein B mRNA editing catalytic polypeptide-like proteins (including APOBEC1, APOBEC2, APOBEC3A, APOBEC3B, APOBEC3C, APOBEC3D, APOBEC3F, APOBEC3G, APOBEC3H, and APOBEC4), recombinant reactive oxygen species modulator 1 (ROMO1), and negative regulator of reactive oxygen species (NRROS). APOBEC1, APOBEC3H gene were lack in the PRJNA656568. In the PRJNA656568, the expression levels of ADAR1, APOBEC3C, APOBEC3D, APOBEC3F, APOBEC3G, and APOBEC4 were higher among SARS-CoV-2 infection than in those not infected by the virus (Figures 6A–6C). In the PRJNA683226, the expression levels of ADARB1, APOBEC1, and APOBEC3H were higher among severe cases than moderate cases. The expression levels of ADARB1 and ADARB2 were higher among severe cases than moderate cases (Figure 6D-F). Further analysis revealed that the expression of APOBEC3A (r = 0.32) was positively correlated with the virus load of SARS-CoV-2 (Figure 6G). However, the expression levels of ADARB2 (r = –0.29), APOBEC2 (r = –0.35), APOBEC4 (r = –0.28), and ROMO1 (r = –0.34) were negatively correlated with the virus load of SARS-CoV-2 (Figures 6H-6K). In the PRJNA631753, these was no significantly difference between SARS-CoV-2 infection and control in the lung (Figures S7A-S7C). The expression levels of ADAR1, ADARB1, APOBEC3C, APOBEC3D, and APOBEC3G were higher in the lung tissues than those in the heart and liver tissues (Figures S5D-S5F). The difference in gender and age was not statistically significant (Figure S6). These results indicated that ADAR and APOBEC family members expression associated with SARS-CoV-2 infection, it might be involved in viral genome mutation.

**Figure 6.**
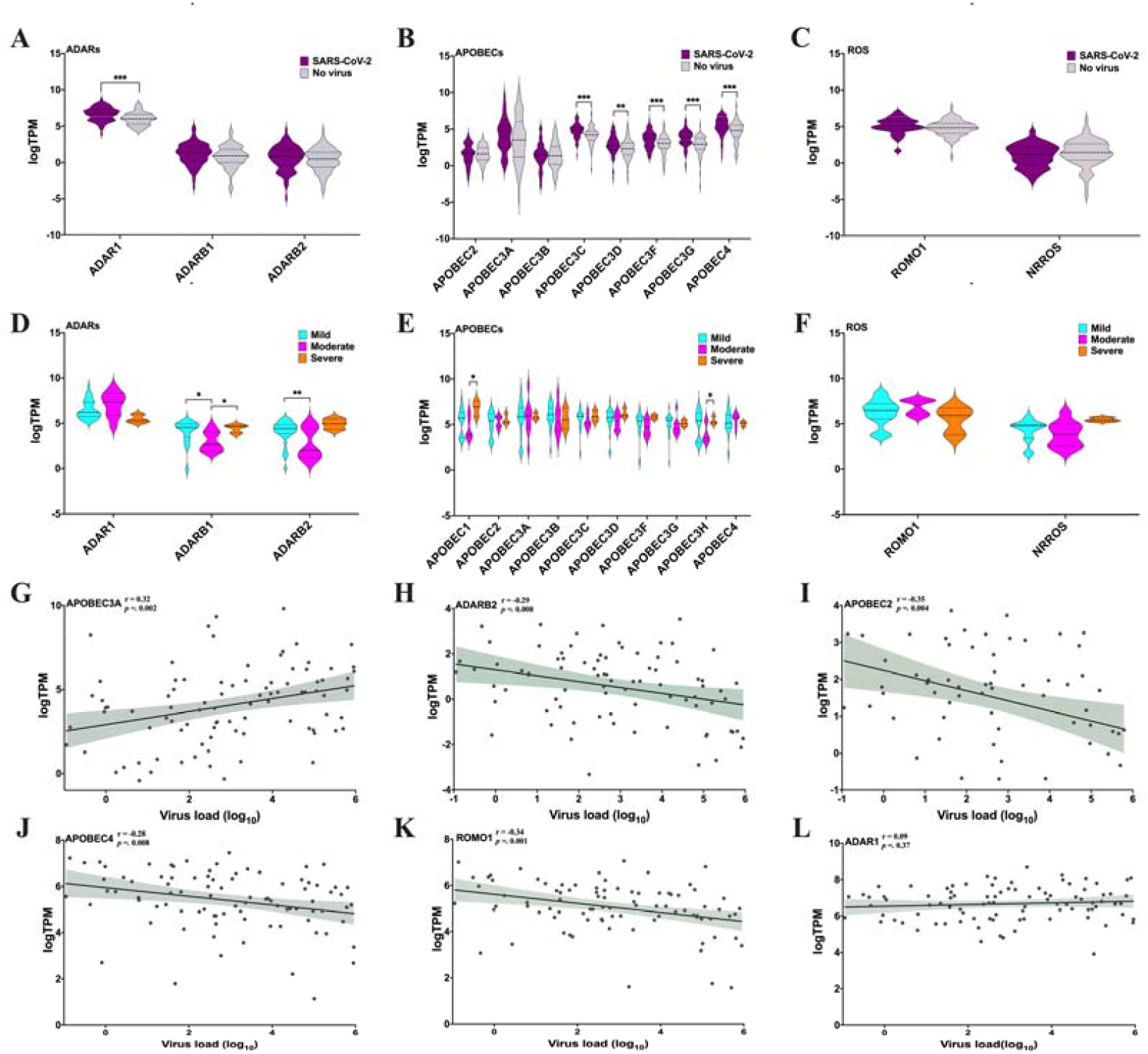
Analysis of expression of RNA-editing enzyme in the transcriptome. A–C: Expression of RNA-editing enzyme after SARS-CoV-2 infection and other infection from nasopharyngeal swabs. D–F: Expression of RNA-editing enzyme in the three clinical outcomes from nasopharyngeal swabs. G–L: Correlation of expression levels of ADARB2, APOBEC2, APOBEC3A, APOBEC4, ROMO1, and NRROS with SARS-CoV-2 virus load. * represents *p* < 0.05, ** represents *p* < 0.01, *** represents *p* < 0.001.

## Discussion

Thousands of mutations in SARS-CoV-2 have been identified. Some of these mutations have become largely fixed in more recent, geographically defined viral populations[8, 25, 26]. Thus, these mutations might be the underlying factors of different clinical outcomes, and they may affect the efficacy of antiviral therapies. Mutations in viral genes may have a direct correlation to clinical outcomes[13]. However, the reason these mutations cause different clinical outcomes remains unknown, a gap in research that requires further analysis. In this study, we identified mutation hotspots in the SARS-CoV-2 sequence associated with clinical outcomes, clades, and regions. We found that the expression levels of D614G, L3930F, V1176F, R203K, G204R, T265I, and Q57H, which were found to be highly distributed in the SARS-CoV-2 VUI 202012/01, 501Y.V2, and 484K.V2 strains, were higher in severe cases than in asymptomatic cases. Moreover, we found that the frequency of nt14408 mutation, which was found in the RdRp gene and was observed to increase the susceptibility to mutations, was higher in severe cases than in asymptomatic cases. Furthermore, we established that the mutation of SARS-CoV-2 was associated with the expression of ADAR and APOBEC.

We identified some mutation hotspots in the SARS-CoV-2 genome that were associated with severe outcomes. The frequency of nt14408 mutation, which was in RdRp (also named nsp12), was higher in severe cases (95.90%) than in asymptomatic cases (56.46%). Notably,the “with nt14408 mutation” group had higher mutation counts than the “without nt14408 mutation” group, especially for the C > T, A > G, G > A, and G > T mutations. However, the mechanism by which nt14408 mutation affects genomic mutation remains unclear. The RdRp is a multi-domain protein that can catalyze the RNA–template dependent formation of phosphodiester bonds between ribonucleotides in the presence of divalent metal ions[27, 28]. In most viruses, RNA polymerase lacks proof-reading capability, with some exceptions such as Nidovirales order (to which the Coronavirus genus belongs), which stands out for having the largest RNA genome. The SARS-CoV-2 RdRp is a key component of the replication/transcription machinery. SARS-CoV-2 shares a higher homology for nsp12 compared with SARS-CoV, suggesting that its function and mechanism of action might be well conserved[29]. Maria Pachetti et al.[30] suggested that the RdRp variant is possibly associated with SARS-CoV-2 mutations. In the present work, the nt14408 mutation in RdRp was found to be more susceptible to mutations, especially for severe outcomes. We speculate that nt14408 mutation may reduce proof-reading capability and make the SARS-CoV-2 genome more susceptible to mutations.

The Q57H variant, which was found to occur in the ORF3a protein, had a higher frequency in severe cases (39.36%) than in asymptomatic cases (10.04%). An RNA-seq dataset (PRJNA631753) from the lungs of deceased patients were also analyzed. Results showed that 85.71% (6/7) of these patients had the Q57H variant. A previous study reported that the Q57H variant can cause dramatic changes in protein structures and decrease the flexibility of domains, thereby enhancing the binding affinities in ORF3a-M and ORF3a-S complexes[31]. Thus, the Q57H variant may destroy drug-targeting sites and lead to therapy failure by shifting the protein-binding interface. Other studies demonstrated that the ORF3a protein can induce apoptosis in cells[32], similar to SARS-CoV [33, 34]. However, SARS-CoV-2 ORF3a has a relatively weaker pro-apoptotic activity than SARS-CoV ORF3a. This property of SARS-CoV-2 ORF3a confers certain advantages: SARS-CoV-2 infection can be relatively mild or even asymptomatic during the early stages, thus allowing the virus to spread more widely. Majumdar et al.[12] reported that the ORF3a mutation is associated with a higher mortality rate in SARS-CoV-2 infection. Therefore, further researche on these mutations is warranted to determine whether these structural alterations in ORF3a influence protein functions and even virus infectivity.

Some of the mutation hotspots identified herein, namely, T265I, D614G, R203K, G204R, L3930F, and V1176F, were found to be associated with severe outcomes with high frequency worldwide. T265I, which is located in non-structural protein 2 (nsp2), has been observed in 13.83% of all cases worldwide. nsp2 is an important domain that ensures the functional integrity of the mitochondria and responds to cellular stress[35]. A change from a polar amino acid (threonine) to a non-polar one (isoleucine) can render nsp2 hydrophobic, thereby inducing structural alterations in that domain[36]. Hence, T265I mutation can change nsp2 structure and mitochondrial permeability, thereby inducing apoptosis pathways. The D614G variant, which was found to occurr in the S protein, has been proved to enhance infection and transmission[9, 11]. The R203K and G204R variants co-occurs in the N protein and cause dramatic changes in protein structure (RMSD ≥ 5.0 Å), thus decreasing the flexibility of the domain[31]. The N protein has a high immunogenic property that enables it to elicit a protective immune response against SARS-CoV-2[37, 38]. R203K and G204R mutations could change N protein structures and affect viral replication, pathogenicity, and disease severity. More importantly, all of these mutation hotspots were also found in the three highly transmissible SARS-CoV-2 variants (i.e., B.1.1.7/VUI202012/01, B.1.351/501Y.V2, and B.1.1.28/484K.V2). These new variants with additional mutation are rapidly spreading in the United Kingdom (VUI 202012/01), South Africa (501Y.V2), and Brazil (484K.V2). These mutation hotspots may be associated with the SARS-CoV-2 pandemic and the resulting mortality. We focus on these mutation hotspots to prevent them from causing the SARS-CoV-2 pandemic again.

Furthermore, we found some mutation hotspots associated with asymptomatic cases. For example, the L3606F mutation in non-structural protein 6 (nsp6) was significantly higher in asymptomatic cases (40.19%) than in symptomatic cases (3.39%). nsp6 mutation could affect viral autophagy[39], a critical host antiviral defense. The issue of whether L3606F mutation can weaken the autophagy function of nsp6 requires further study. Notably, these major mutations in asymptomatic cases mostly co-occurred and were distributed in different genes. For example, R6997P (ORF1Ab), A222V (S), A220V (N), and V30L (ORF10) co-occurred in 0.5% of all cases in July 2020 and exceeded 50% of all cases by January 2021 worldwide. M3087I (ORF1Ab), K4576N (ORF1Ab), X5167Y (ORF1Ab), N5542D (ORF1Ab), S5585I (ORF1Ab), and A376T (N) co-occurred in 0.8% of all cases in January 2020 to 5.7% of known cases in January 2021 worldwide. Thus, the mutation is a cumulative process. This significant result indicated that co-occurring mutations are an important factor for the SARS-CoV-2 pandemic. These mutations may be relevant in designing vaccines.

The factors that caused these different mutations are unclear. The process of viral genome mutagenesis includes host-dependent RNA editing enzymes, RdRp, spontaneous nucleic acid damages due to physical and chemical mutagens and recombination events. The host-dependent RNA editing enzymes includes ADAR, APOBEC, and reactive oxygen species (ROS) (regulated by ROMO1 and NRROS). A previous study offered a possible evidence for host-dependent RNA editing in the transcriptome of SARS-CoV-2[40]. Here, we analyzed varous SNV types and host editing enzymes. The SNV types were dominated by C>T, A>G, and G>T. The C>T substitution is a signature likely related to APOBEC activity. The A>G substitution is probably related to ADAR activity. The G>T substitution is possibly related to ROS-related processes. The expression levels of ADAR1, APOBEC3C, APOBEC3D, APOBEC3F, APOBEC3G, and APOBEC4 increased after SARS-CoV-2 infection, indicating that ADARs and APOBECs are involved in SARS-CoV-2 infection. ADAR, which can convert A-to-I, has three ADAR genes encoded in the human genome[41]. APOBEC, a family of zinc-dependent deaminases, includes APOBEC1, APOBEC2, AID, APOBEC3 (with family members A, B, C, D, F, G, and H), and APOBEC4[42]. Most APOBECs have been shown to catalyze cytosine deamination to uracil (C-to-U) of foreign single-stranded DNA and RNA[43]. We found that clinical outcomes could affect the expression of ADAR and APOBEC, a process that is probably the main cause of different mutation rates among clinical outcomes. Notably, the expression levels of ADAR3, APOBEC2, APOBEC4, and ROMO1 were negatively correlated with the virus load of SARS-CoV-2, indicating that these deaminases may be related to antivirus.

However, this research has several limitations. First, the insufficient clinical information on these sequences may have led to missing some mutations associated with clinical outcomes. Second, given that some raw RNA-seq data were unavailable, the relationship between SARS-CoV-2 mutation and clinical outcomes at the patient level was not analyzed. Third, some hotspot mutations lacked experimental proof and thus require experimental verification.

## Conclusion

By analyzing 209,551 high-coverage complete virus sequences and 352 RNA-seq samples, we found that several hotspot variants, especially Q57H, T265I, D614G, R203K, and G204R mutations, were associated with severe outcomes. These mutations were higher in severe cases than in asymptomatic cases and were highly distributed in the SARS-CoV-2 VUI 202012/01, 501Y.V2, and B.1.1.248 strains. The nt14408 mutation located in RdRp was accompanied with greater susceptibility to mutations in the SARS-CoV-2 genome. We also found that the main SNV types of SARS-CoV-2 were C>T, A >G, and G >T, all of which were associated with the expression levels of ADARS and APOBEC. Our results indicated that these variants must be further investigated. This study provides insights into the development of diagnostic and therapeutic strategies for COVID-19.

## Supporting information

Fiure S1

## Acknowledgements

We are grateful to Dr. Huifeng W (Guangxi Medical University) for aiding us in drawing the figures. We express our appreciation to Chenyang H and Tingxi L for assisting us with the data analysis. This study was funded by Guangxi Key Research and Development Program (No: Guike AB20059002, GuikeAB20072005).

## Author contributions

L.H.Y. J.L. and W.X.P. conceived the study. L.P. designed the analysis code. S.F.Z. L.S.Q. Q.H.W. Y.F.W. B.X. X.X. L.X.L. C.Y.Y. L.C.W analyzed the data. L.H.Y. and W.X.P. wrote the manuscript. All authors have reviewed the manuscript.

## Data availability

All data are provided in the article and supplementary information. The data may also be obtained from the corresponding author upon reasonable request. Source data are provided with this paper.

## Code availability

All data analysis codes are available from the corresponding author upon reasonable request.

## Supplementary Information

**Figure S1** Analysis of the factors associated with SNV types. A: SNV types distributed in different clades; B: SNV types distributed in different regions. C: SNV types distributed in different ages; D: SNV types distributed in different genders. ^a^ *p* < 0.001 represents there is significant difference among the eight clades. ^b^ *p* < 0.001 represents there is significant difference among the six regions.

**Figure S2** Motif of different SNV types in SARS-CoV-2, SARS and MERS. A: Motif of different SNV types in SARS-CoV-2; B: Motif of different SNV types in SARS; C: Motif of different SNV types in MERS.

**Figure S3** Motif of different SNV types in SARS-CoV-2 among different clinical outcomes. A: Motif of A>G mutation among different clinical outcomes; B: Motif of C>T mutation among different clinical outcomes; C: Motif of G>A mutation among different clinical outcomes; D: Motif of G>T mutation among different clinical outcomes.

**Figure S4** Frequency of all mutations among different clinical outcomes. A: Frequency of all mutations among asymptomatic cases. B: Frequency of all mutations among mild/moderate cases. C: Frequency of all mutations among severe/ICU/deceased cases.

**Figure S5** Distribution of all SNV types among different samples.

**Figure S6** Expression of ADARs, APOBECs, and ROS among gender and age. A–C: Expression of ADARs, APOBECs, and ROS among genders. D–F: Expression of ADARs, APOBECs, and ROS among ages.

**Figure S7** Expression of ADARs, APOBECs, and ROS in different tissues. A–C: Expression of ADARs, APOBECs, and ROS in lung tissues after SARS-CoV-2 infection. D–F: Comparison of the expression levels of ADARs, APOBECs, and ROS in lung, liver, and heart tissues.

