## Supplementary figures and images for "Emerging SARS-CoV-2 mutation hotspots associated with clinical outcomes"

### Figure S2

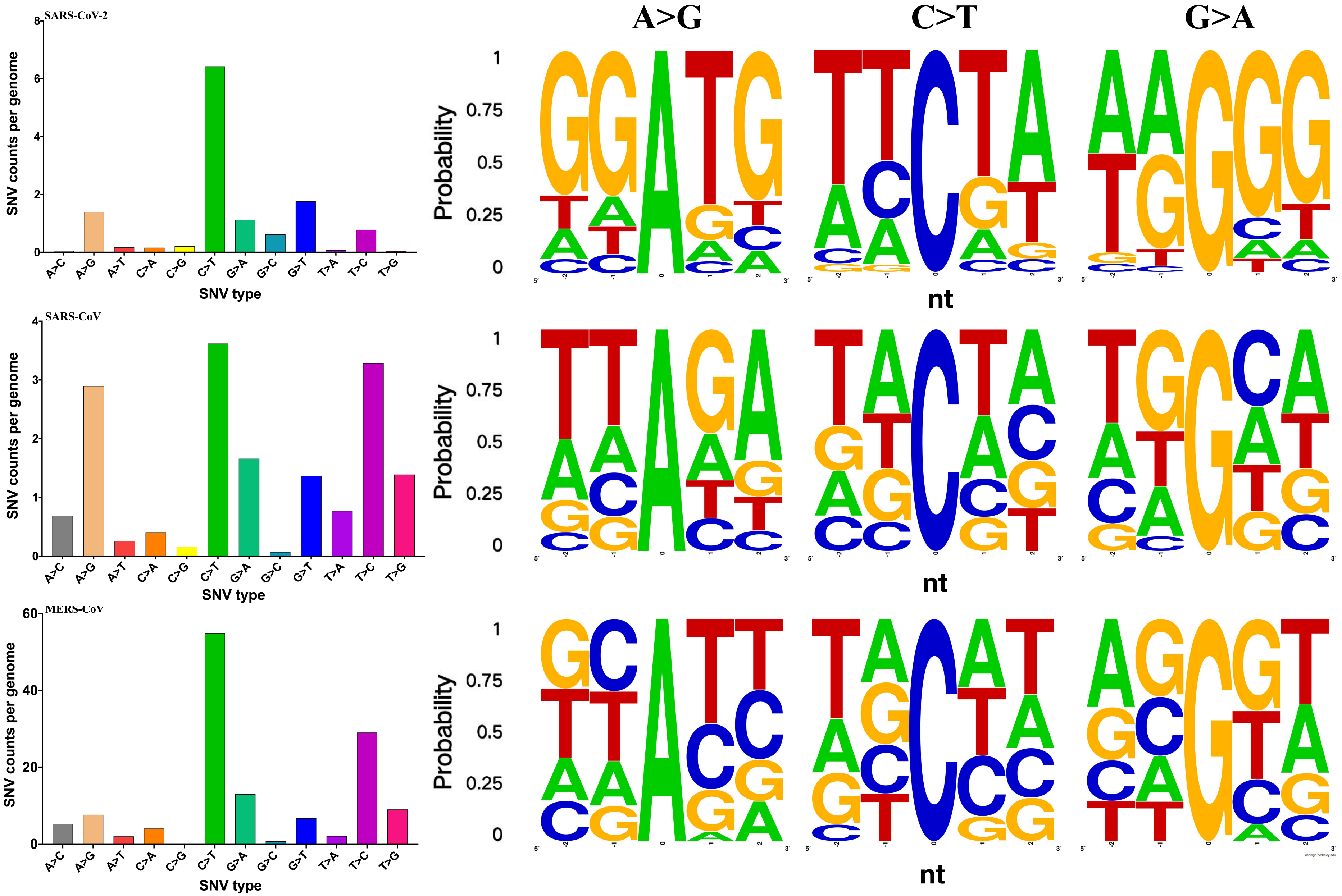

### Figure S3

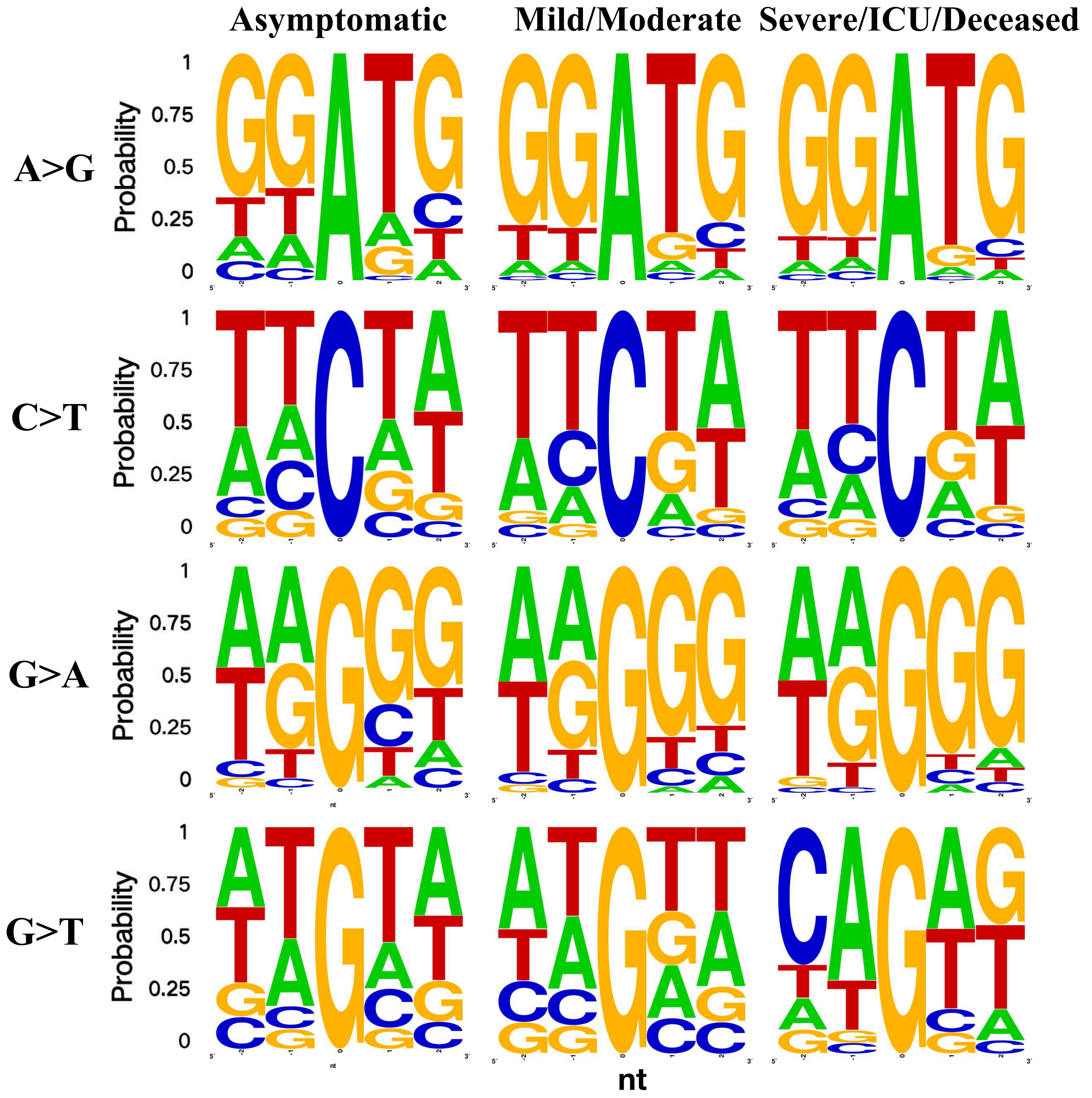

### Figure S4

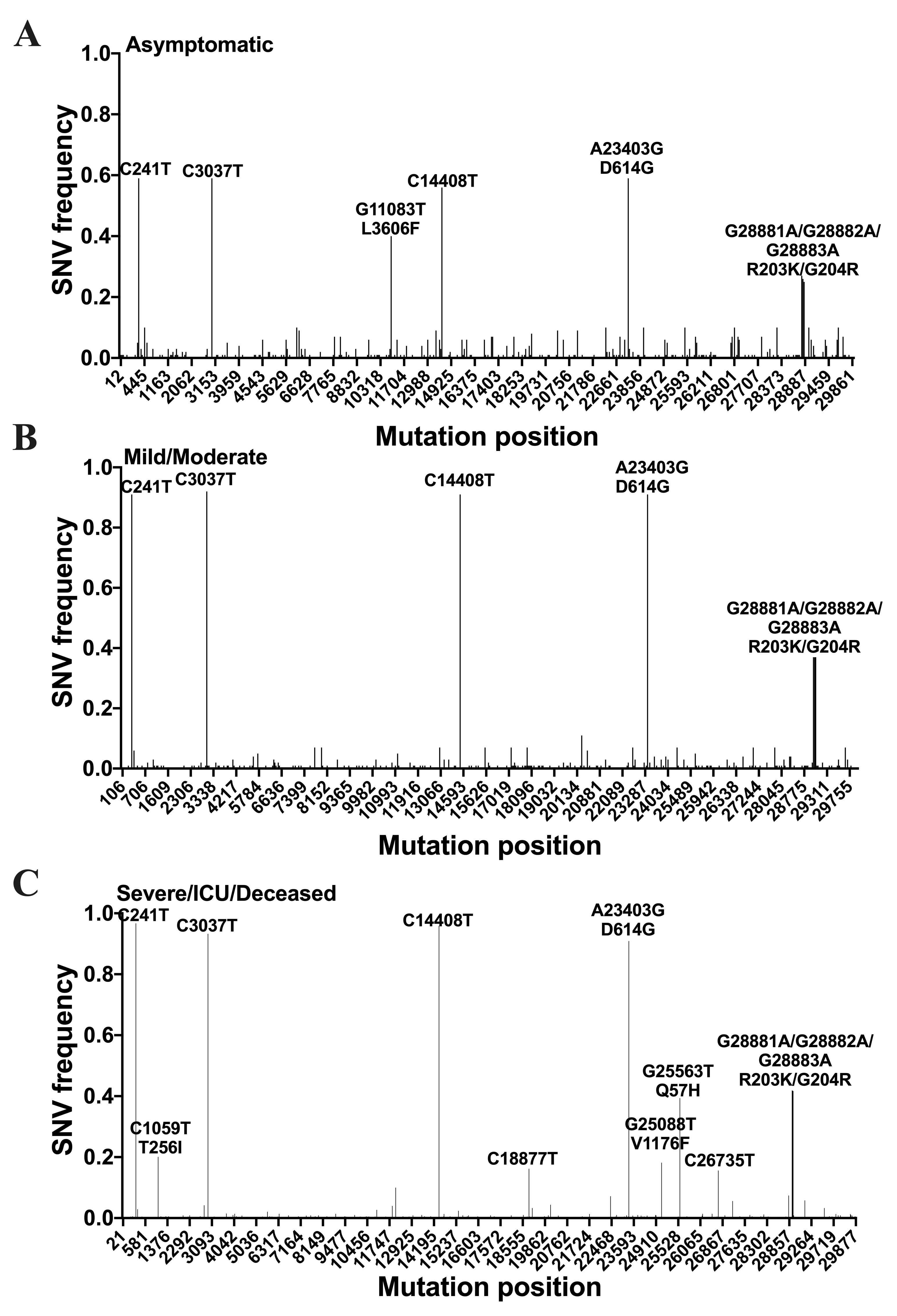

### Figure S5

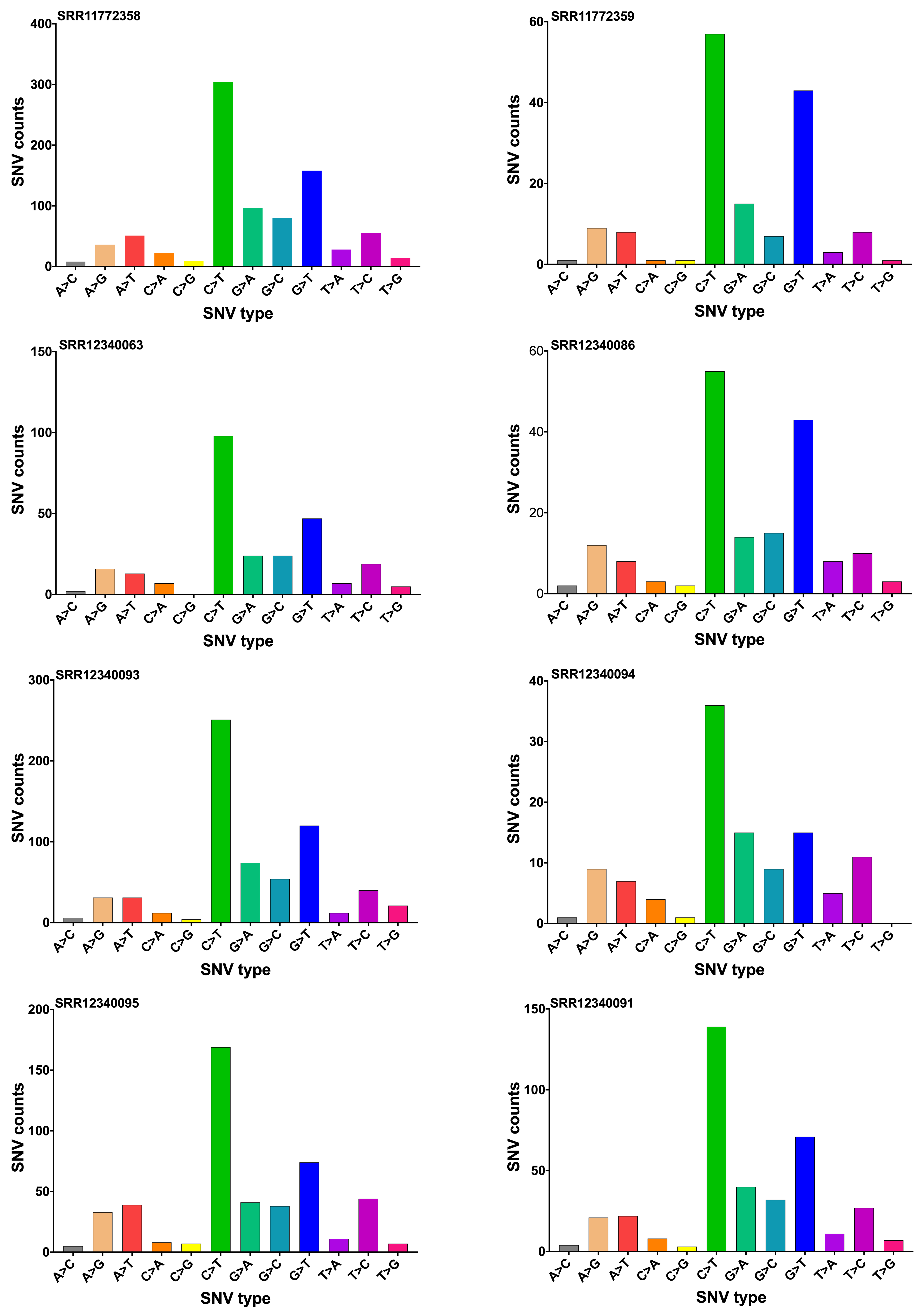

### Figure S6

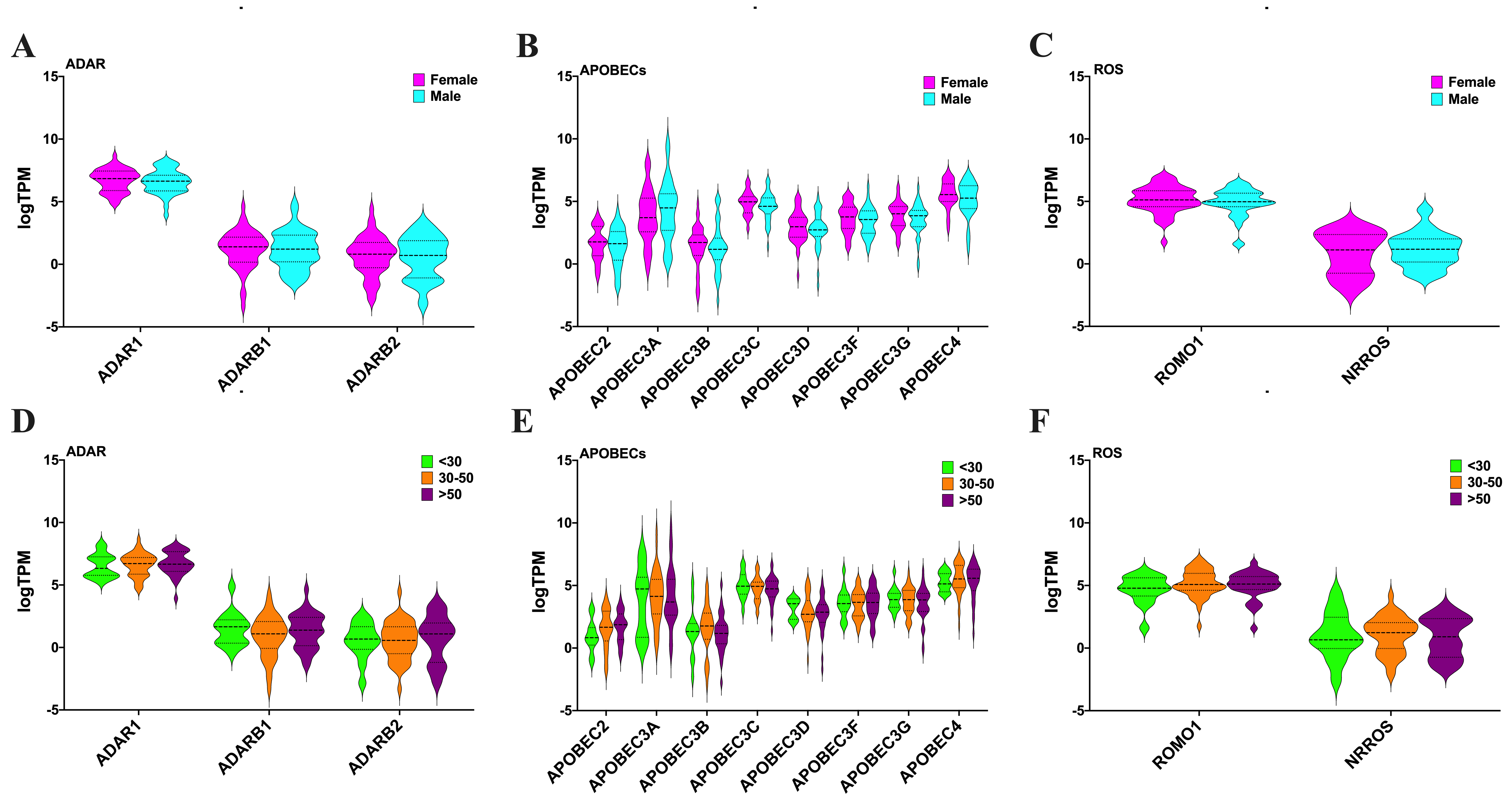

### Figure S7

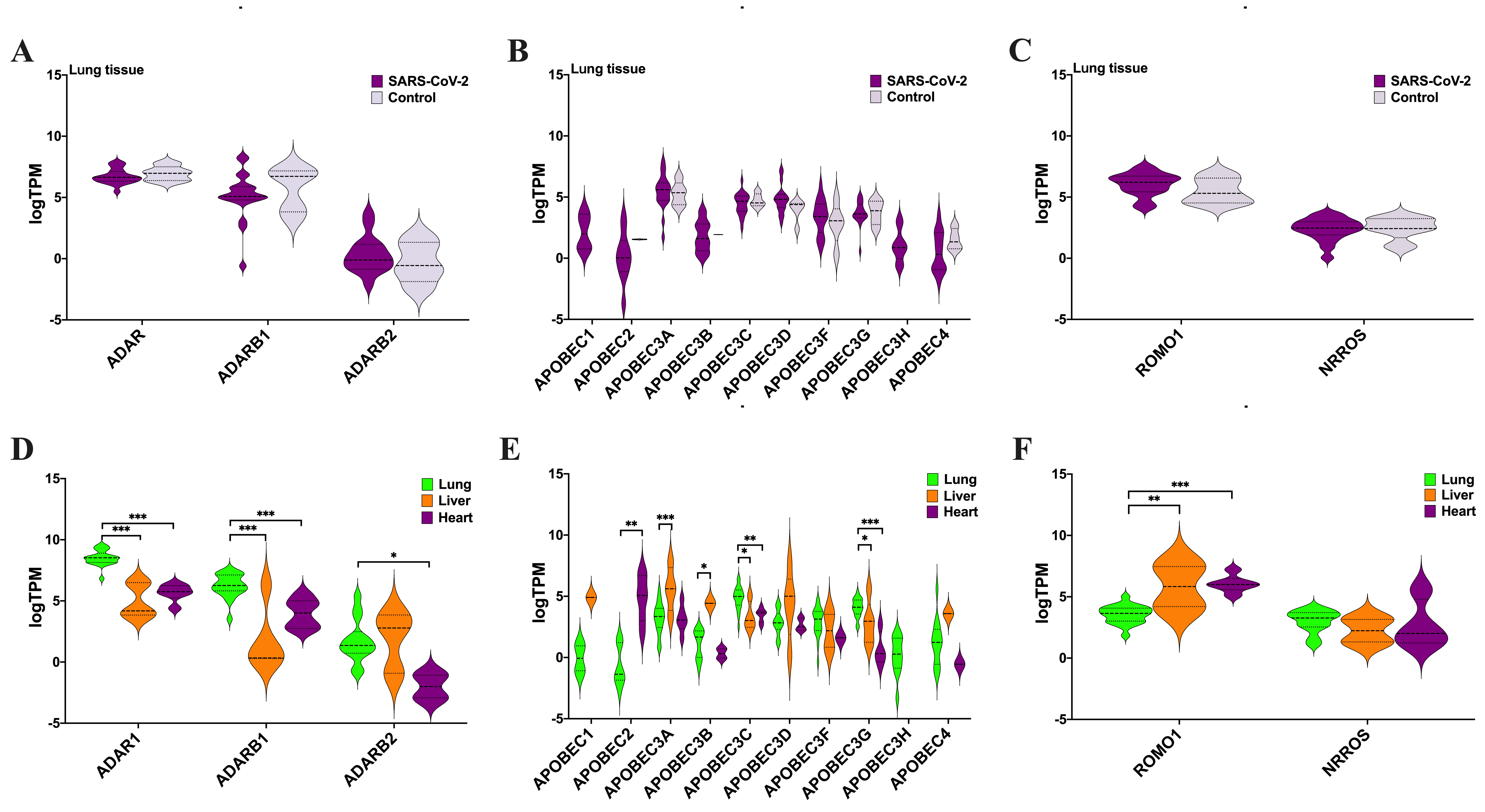

### Fiure S1

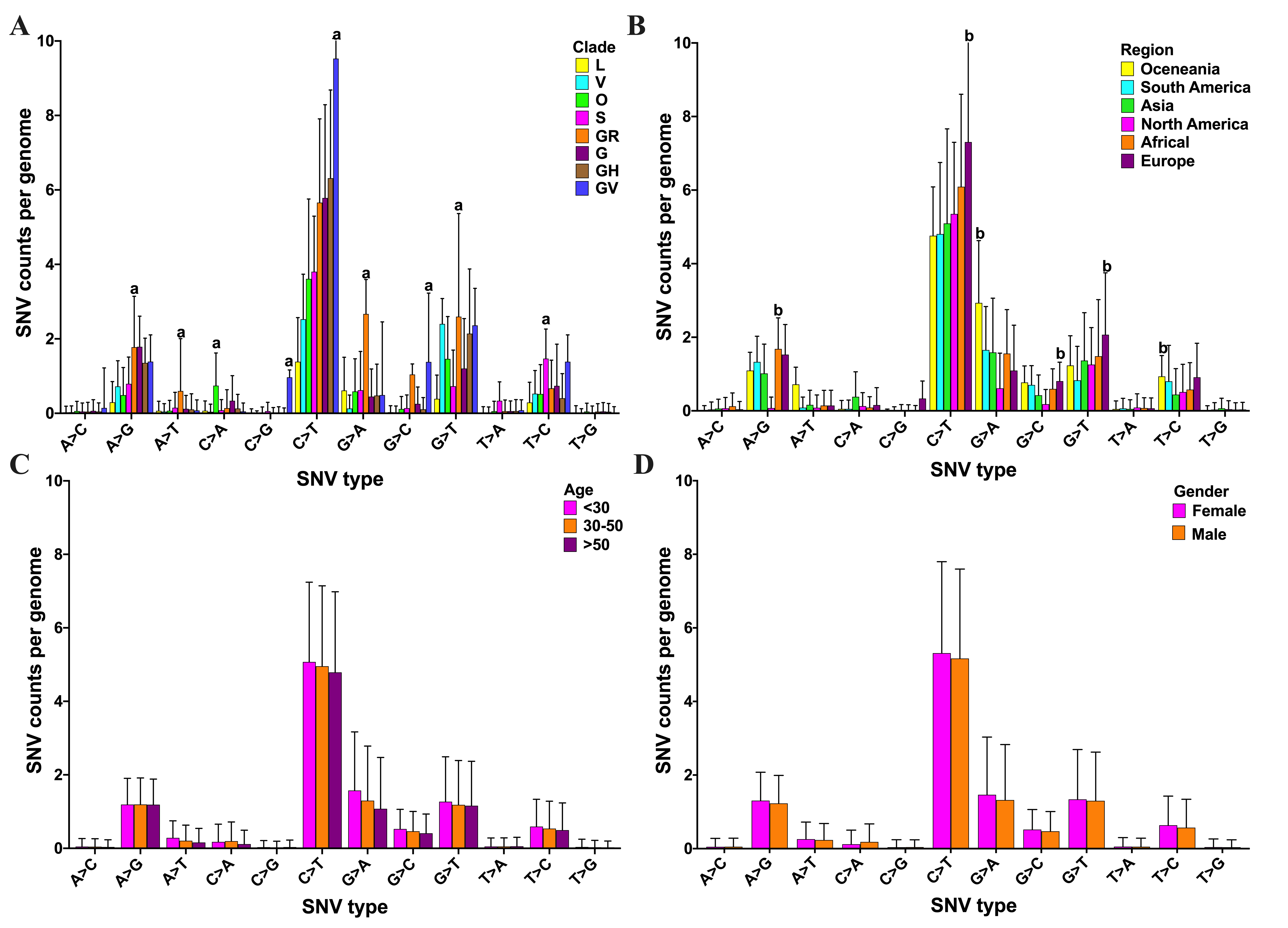
